# Generic solving of one-compartment toxicokinetic models

**DOI:** 10.1101/2021.05.06.442956

**Authors:** Sandrine Charles, Aude Ratier, Christelle Lopes

**Affiliations:** University of Lyon; University Lyon 1; CNRS; UMR 5558, 43 boulevard du 11 novembre 1918, Villeurbanne Cedex, F-69622, France

**Author notes:** Corresponding author *Email address:* (Sandrine Charles).

**Keywords:** ordinary differential equations, environmental risk assessment, living organisms, active substances, TK models, bioaccumulation metrics

## Abstract

This paper gives the full analytical solution of the generic set of ordinary differential equations that define one-compartment toxicokinetic models. These models describe uptake and elimination processes taking place within living organisms when exposed to chemical substances. The models solved in this paper consider living organisms as a unique compartment, into which a parent compound enters via several possible exposure routes and from which it is eliminated as well as its potential metabolites. Benefiting from generic solutions of one-compartment toxicokinetic models is particularly useful when fitting them to experimental data, facilitating the writing of the inference algorithms leading to parameter estimates. Additionally, these models are of crucial interest in environmental risk assessment for the calculation of bioaccumulation metrics as required by regulators in support of decision making when they evaluate dossiers for marketing authorisation of active substances.

Graphical abstract

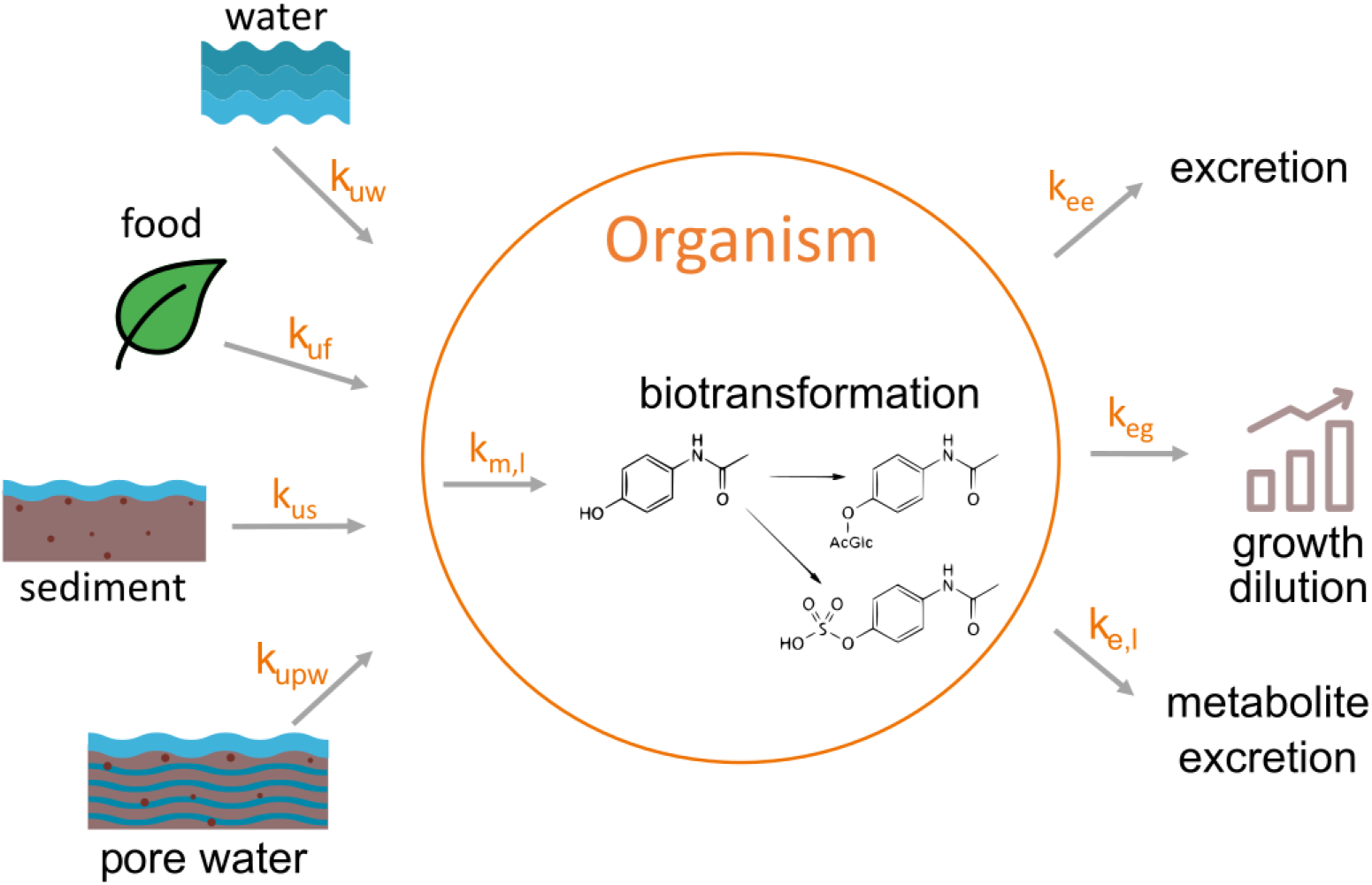

## 1. Introduction

In this paper, we consider a very generic one-compartment toxicokinetic (TK) model describing uptake and elimination processes taking place within living organisms when exposed to chemical substances. On a general point of view, the term TK refers to the movement and fate (also called disposition) of chemical substances, potentially toxic. In particular, this term is used when describing the time course of absorption, distribution and elimination (accounting for both bio-transformation and excretion) of substances within organisms (ADME). Toxicokinetics is also related to pharmacokinetics, even sometimes considered as the same scientific field, the main difference being linked to the type of compound (toxicants or pharmaceuticals). In addition, in comparison with pharmaceutical exposure, exposure to toxicants is often uncontrolled and variable over time, also at larger concentrations [10].

Mathematical TK models are particularly useful when it is needed to predict adverse effects of xenobiotics or to prevent undesired residues within animal tissues from entering the human food chain [4]. Indeed, TK models allow to predict tissue concentrations over time, involving several kinetic parameters, such as uptake and elimination rates, from which useful metrics for assessing risk may be derived [19]. But TK models can also be used in support of a better understanding of the underlying physiological mechanisms driving chemical ADME processes [13, 12].

A wide variety of TK models exists, chosen from both the available data and the model’s purpose [23]. Basically, we distinguish classic TK models from physiologically-based TK models, referred to as PBTK, or PBPK models in the filed of pharmacology. As fully detailled in [7], PBPK models are a crucial tool in Environmental Risk Assessment (ERA) for regulatory bodies such as the US EPA (US Environmental Protection Agency) or the EFSA (European Food Safety Authority) [16, 15]. In fact, PBPK models are an ethical and scientifically sound method to predict exposure to toxic xenobiotics in humans through animal-to-human extrapolation or based on human biomonitoring data. But (PB)TK models are also of particular interest in ERA, when the calculation of bioaccumulation metrics is required by regulators in support of decision making in evaluating dossiers for substance marketing authorisations [18, 17, 3].

The generic TK model we unravel in this paper is a one-compartment model considering organisms as a whole in which chemical compounds may enter and from which these compounds can be eliminated). More complex TK models (namely PBTK models) refine the description of contamination pathways within organisms, distinguishing organs and tissues from physiological hypotheses on potential targets of exposure compounds; PBTK models have been mainly developed for humans [25, 24] or big mammals [9, 14]; a very recent PBTK model has been proposed by [8] for invertebrates, pushing the immense potential of these models a step further.

Our TK model, even if not fully refined as could be a PBTK model, has the great advantage of accounting for several exposure sources (*e.g.*, by water, sediment and/or food), several elimination processes (*e.g.*, direct elimination, dilution by growth and/or biotransformation) and several potential metabolites of the parent chemical compound to which organisms are exposed [21]. The next section is a fully detailed description of the dynamical system of ordinary differential equation ( ODE) and its mathematical exact solution, with a summary in section 3. Then section 4 illustrate the use of model, based on simulations from parameter values obtained by the study of real data sets.

## 2. The one-compartment TK set of ODE

Our generic one-compartment TK model is composed in total of two sets of ODE, one set for the accumulation phase (that includes both absorption and elimination processes, as those related to the biotransformation of the parent compound into metabolites) and during which organisms are exposed to a given compound; the second one for the depuration phase (with only elimination processes including biotransformation) and during which organisms are transferred into a clean medium. The transition from one set of ODE to the other takes place at time *t_c_* corresponding to the duration of the accumulation phase (see Table 1).

**Table 1:**
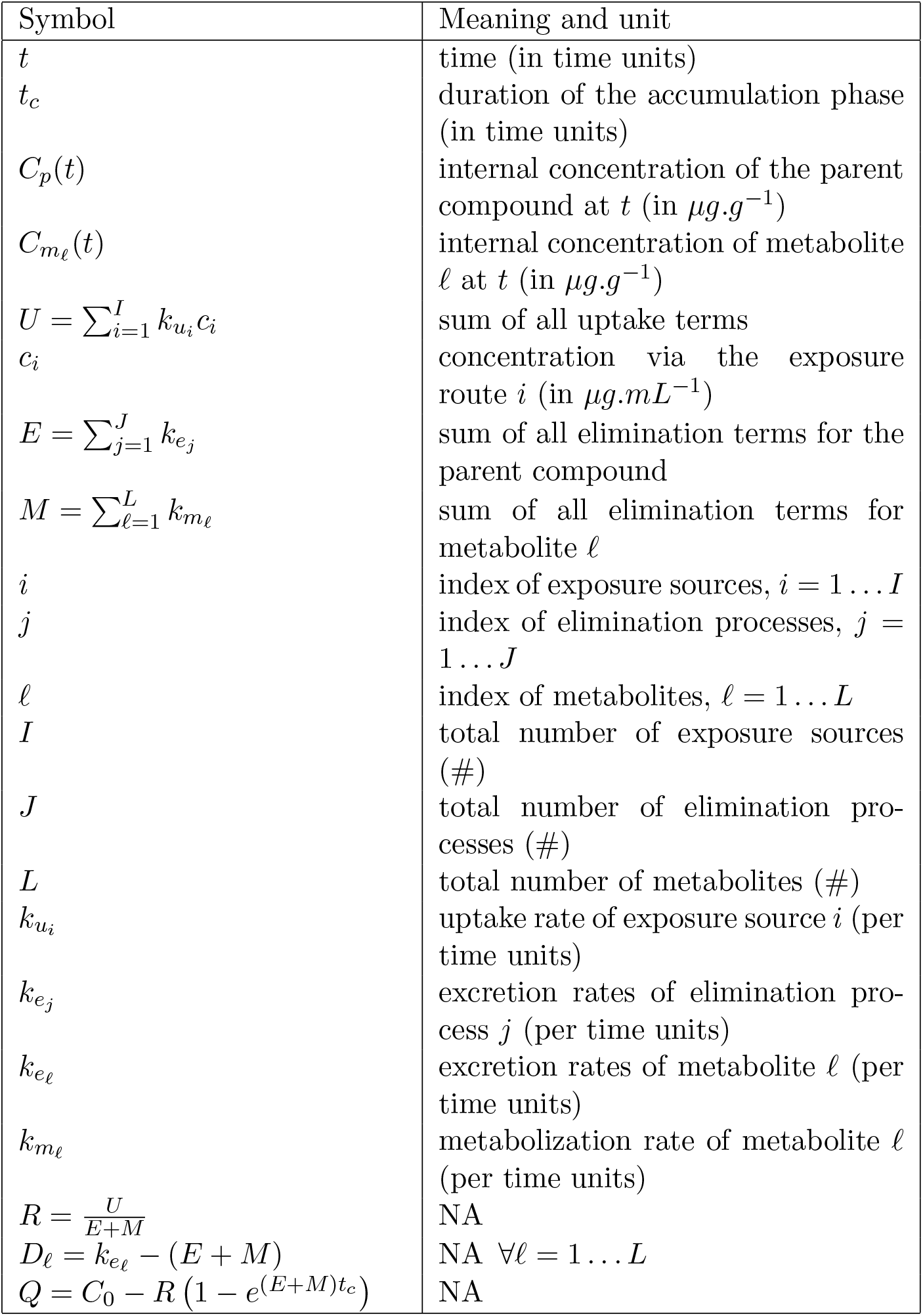
Symbols, meaning and units for all parameters and variables involved in the full set of ordinary differential equations defining the generic on e-compartment toxicokinetic model. # stands for numbers, while NA means ‘Not Applicable’.

In practice, the exposure concentration to which organisms are exposed may vary over time as in real environments, but then, there is no analytical solution of the TK model; only a numerical solution can be obtained with an appropriate algorithm. Our paper thus assumes that the exposure concentration remains constant over time whatever the exposure source. Such an experimental condition can be ensured for most of the chemical compound when performing laboratory experiments. In addition, this assumption allows to provide the exact solution of the full one-compartment TK model by considering as many routes of exposure and as many elimination processes as desired, as well as an infinite number of phase I metabolites, *i.e.*, directly derived from the parent compound to which organisms are exposed. Note that [11] already provided a partially resolved one-compartment TK model but only for the accumulation phase and one exposure route. In a similar manner, [6] published an analytical solution for a linear multi-compartments TK model wit a non-zero initial condition.

### 2.1. Accumulation phase (0 ≤ t ≤ t_c_)

The set of ODE describing the accumulation phase (0 ≤ *t* ≤ *t_c_*) writes as follows:

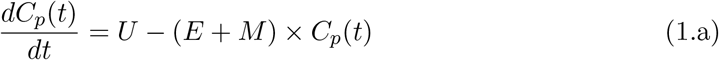

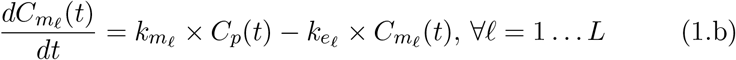

All parameters and variables, with their meaning and units when applicable, are gathered together in Table 1.

Equation (1.a) for the parent compound is a linear first-order ODE with constant coefficients and a second member. Equation (1.a) admits 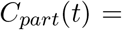 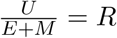 as a particular solution. Equation (1.a) without its second member writes:

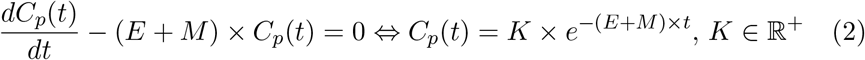

Given the initial condition *C*(*t* = 0) = *C*_0_ (*C*_0_ ≥ 0), we finally get the full analytical solution of Equation (1.a), providing the internal concentration of the parent compound over time during the accumulation phase (0 ≤ *t* ≤ *t_c_*) as follows:

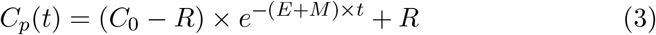

See Table 1 for the definition of parameter *R*.

Equation (1.b) is also a linear first-order ODE with constant coefficients and a second member, with the following analytical solution when removing its second member:

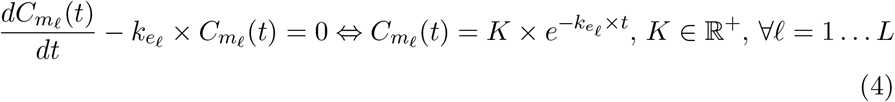

The method of variation of a constant consists of writing the general solution of Equation (1.b) as:

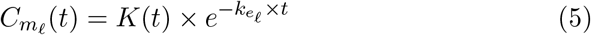

and to find function *K*(*t*) by deriving 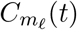 and re-injecting the result into Equation (5). The derivative from Equation (1.b) writes:

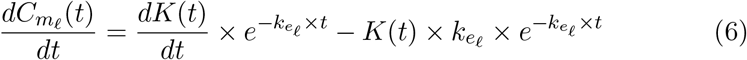

while the re-injection into Equation (1.b) leads to:

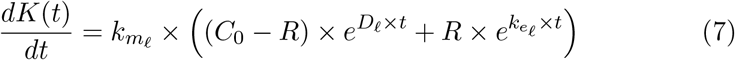

which integrates into:

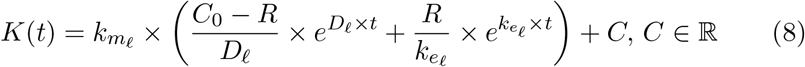

See Table 1 for the definition of parameter *D_l_*.

The general solution of Equation (1.b) finally writes as follows:

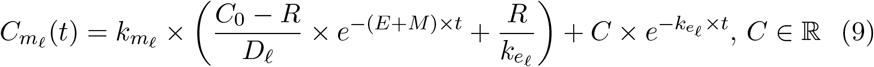

When considering the following initial condition 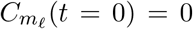, biologi cally meaning that when the accumulation phase starts, organisms are only exposed to the parent compound, so that there are no metabolites within. Then, we finally get the full analytical solution of Equation (1.b) providing the internal concentration of metabolite *l* over time during the accumulation phase:

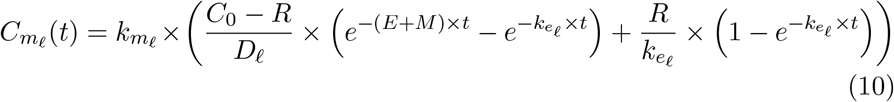

### 2.2. Depuration phase (t ≥ t_c_)

The set of ODE describing the depuration phase (*t ≥ t_c_*) writes as follows:

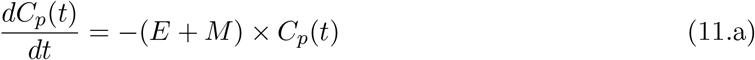

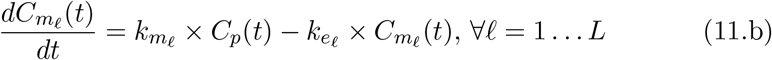

All parameters and variables, with their meaning and units when applicable, are gathered together in Table 1.

Equation (11.a) is a linear first-order ODE without a second member, so that it has a general solution of the following form:

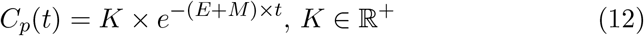

For the depuration phase and the parent compound, the initial condition comes from the calculation of the internal parent compound concentration at the end of the accumulation phase (*i.e.*, at time *t = t_c_*) thanks to solution (3):

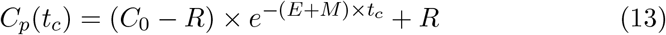

From the general analytical solution given by Equation (12), we get *C_p_*(*t_c_*) = *Ke*^−(*E*+*M*)*t_c_*^ leading to a constant *K* in Equation (12) equals to:

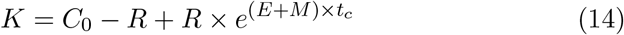

Then, the final analytical solution of Equation (11.a) providing the internal concentration of the parent compound over time during the depuration phase (*t ≥ t_c_*) writes:

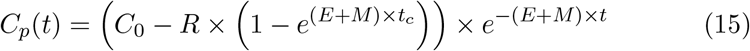

For simplicity reasons, Equation (15) above can be written as *C_p_*(*t*) = *Qe*^−(*E*+*M*)×*t*^ with *Q* as defined in Table 1.

Equation (11.b) is a linear first-order ODE with constant coefficients and a second member. The analytical solution of Equation (11.b) without its second member writes:

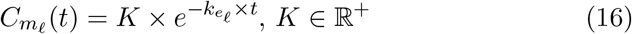

As previously, the method of the variation of a constant provides the general solution of Equation (11.b) as 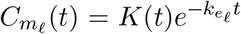, requiring to search for function *K*(*t*).

The derivative of 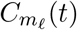 from Equation (16) writes:

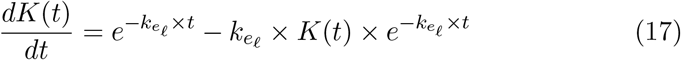

The re-injection of derivative (17) into Equation (11.b) leads to:

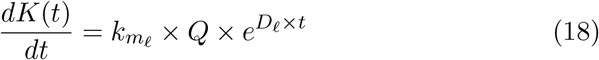

which integrates into:

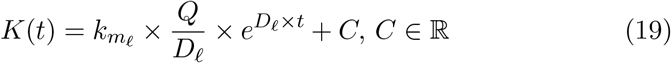

finally leading to the general analytical solution of Equation (11.b):

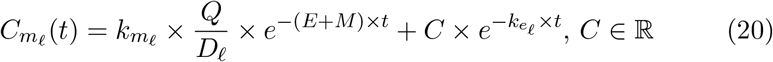

Constant *C* can be determined from the initial condition, *i.e*., from the internal concentration of metabolite *l* at *t = t_c_* both at the end of the accumulation phase and at the beginning of the depuration phase. From Equation (20), we get 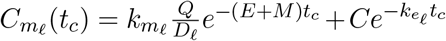, and from Equation (10), we get 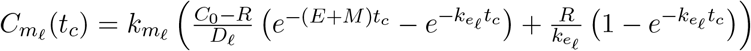. Finally, we get the following expression for constant *C*:

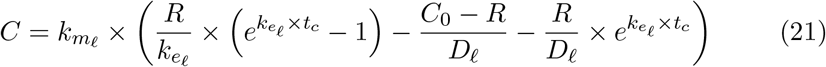

Replacing constant *C* in Equation (20) gives the final analytical solution of Equation (11.b) providing the internal concentration of metabolite *l* for the depuration phase (*t > t_c_*) as follows:

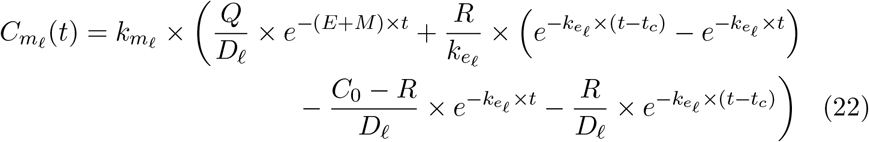

Replacing constant *Q* by its own expression (Table 1) leads to:

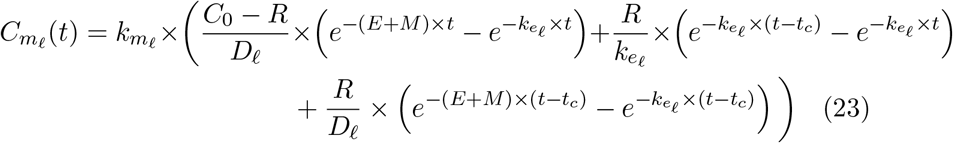

## 3. The generic set of solutions

Remembering the following intermediate notations 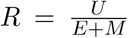 and 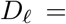 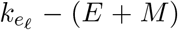 we finally obtain the full set of analytical solutions corresponding the whole one-compartment TK set of ODE describing the time course for both the accumulation and the depuration phases of the parent compound and its potential metabolites.

- The analytical solution for the internal concentration of the parent compound during the accumulation phase, previously referred as Equation (3):

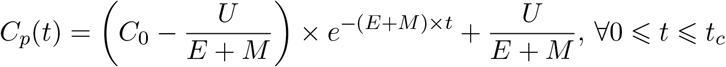
- The analytical solution for the internal concentration of metabolite *l* during the accumulation phase, previously referred as Equation (10):

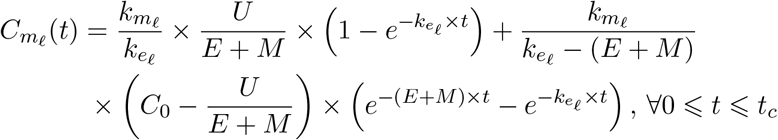
- The analytical solution for the internal concentration of the parent compound during the depuration phase, previously referred as Equation (15):

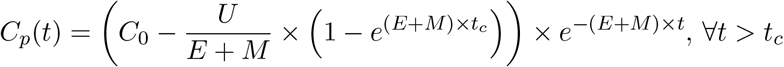
- The analytical solution for the internal concentration of metabolite *l* during the depuration phase, previously referred as Equation (23):

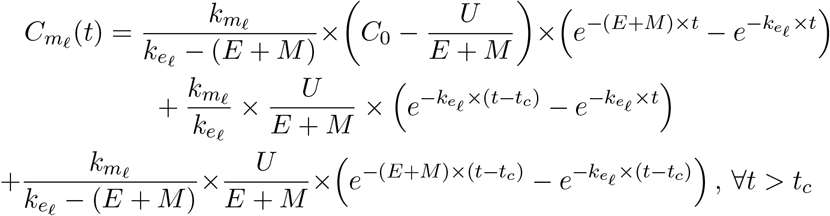

We could fully finish the writing of the very final generic analytical solution of the one-compartment TK model with all the parameters to estimate from observed data using an inference process by replacing constants *U*, *E* and *M* by 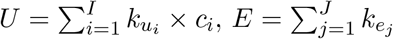 and 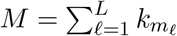, respectively.

## 4. Model simulations based on real case-studies

Today, the above exact mathematical solution of the generic one-compartment TK model is used in support of the development of the R-package `rbioacc` [2], as well as its web interface MOSAIC*_bioacc_* freely available at https://mosaic.univ-lyon1.fr/bioacc [22]. Both tools allow fitting the TK model to bioaccumulation data in a user-friendly way, providing all necessary outputs to check for goodness-of-fit and obtain bioaccumulation metrics, whatever the considered species-compound combination. Associated to the web interface, a database is also made available on-line, http://lbbe-shiny.univ-lyon1.fr/mosaic-bioacc/data/database/TK_database.html, with a wide collection of accumulation-depuration data in support of toxicokinetic modelling [20]. From this database, we selected three real case-studies from which we got fitting results (especially the joint posterior probability distribution of all kinetic parameters) in order to perform the subsequent model simulations, thus illustrating the use of the generic TK model.

All model simulations of the generic set of mathematical solutions (Equations (3), (10), (15) and (23)) were performed under the R software with the `rbioacc` package and function `predict()` [2]; note that the same simulations can be performed directly online with the platform MOSAIC*_bioacc_* and its prediction module (see here for the prototype version: https://scharles-univlyon1.shinyapps.io/bioacc-tabs). We used 500 time points in [0; *t_f_*] where *t_f_* stands for the final time of each simulation. These simulations illustrate the toxicokinetic in three case studies where different compounds are bioaccumulated by different species. In order to proceed, the time duration of the accumulation phase (parameter *t_c_*) is required, as well as the exposure concentrations in the media and the model parameter values (Table 2); all associated numerical values were taken from fitting results performed with real data sets. Figures 1 to 5 show the simulations of internal concentrations over time for the three species-compound combinations fully described hereafter. For each case study, TK model parameters were varied one-at-a-time with 20, 50 and 80% of increase from their original value.

**Table 2:** Combinations of inputs for model simulations according to Figure 1. Hyphens stands for no required inputs.

| Input | Units | Simple<br>(section 4.1) | Multiple exposures<br>(section 4.2) | Biotransformation<br>(section 4.3) |
| --- | --- | --- | --- | --- |
| $c_w$ | $\mu g.mL^{-1}$ | 0.0044 | - | 15.53 |
| $c_s$ | $\mu g.g^{-1}$ | - | 56.6 | - |
| $c_f$ | $\mu g.g^{-1}$ | - | 1.46 | - |
| $t_c$ | *days* | 49 | 7 | 1 |
| $k_{uw}$ | *days*<sup>-1</sup> | 10.46 | - | 16740 |
| $k_{us}$ | *days*<sup>-1</sup> | - | 0.071 | - |
| $k_{uf}$ | *days*<sup>-1</sup> | - | 0.013 | - |
| $k_{m1}$ | *days*<sup>-1</sup> | - | - | 73.27 |
| $k_{m2}$ | *days*<sup>-1</sup> | - | - | 0.5166 |
| $k_{m3}$ | *days*<sup>-1</sup> | - | - | 0.1957 |
| $k_{ee}$ | *days*<sup>-1</sup> | 0.04 | 0.178 | 4.164 |
| $k_{e1}$ | *days*<sup>-1</sup> | - | - | 561 |
| $k_{e2}$ | *days*<sup>-1</sup> | - | - | 0.123 |
| $k_{e3}$ | *days*<sup>-1</sup> | - | - | 0.7808 |

**Figure 1:**
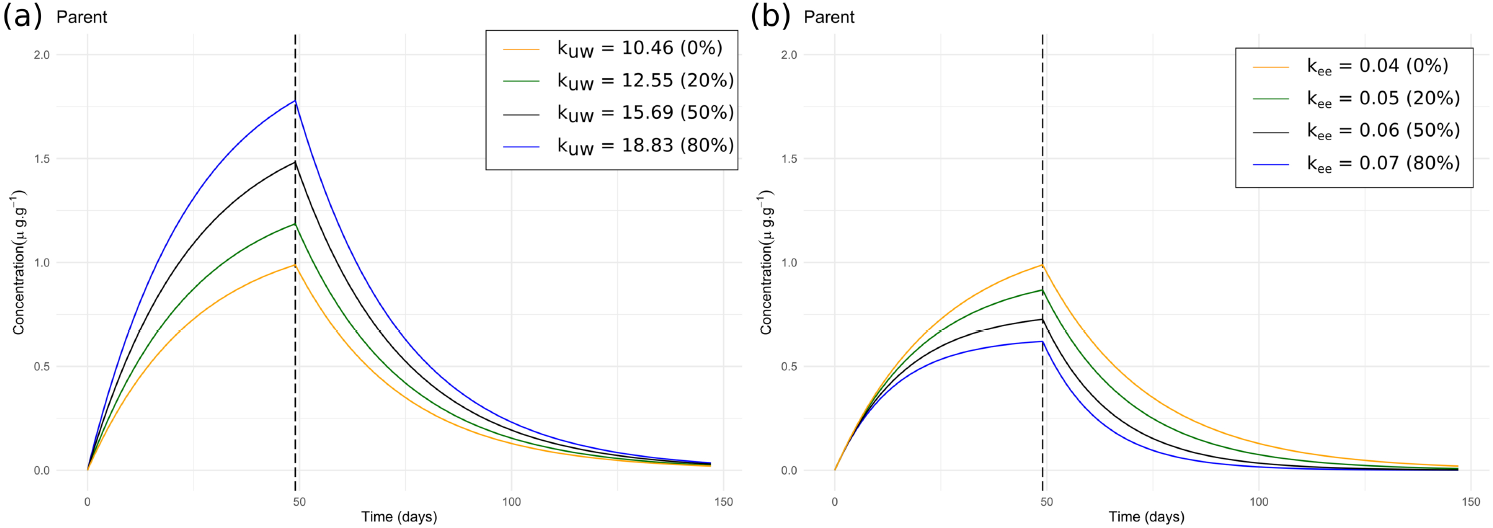
Example of model simulations for a simple TK model (exposure by water and elimination by excretion) and the influence of variations of parameters (a) 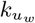 (uptake rate) and (b) 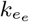 (excretion rate).

### 4.1. The most elementary one-compartment TK model

Equations (3) and (15) are simulated for a simple case study where fish are exposed to a highly hydrophobic chemical contaminated water. Only excretion is considered [5] (Figure 1 and Table 2). The corresponding inputs are the exposure concentration *c_w_* (referring to *c_i_* and *I* = 1), the uptake rate from water 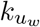 (referring to 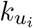 and *I* = 1) and the excretion rate 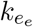 (referring to 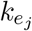 and *J* = 1). When parameter 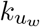 increases, internal concentrations are higher (*e.g.*, blue curve in Figure 1.a) than with the original value (orange curve in Figure1.a). Biologically speaking, the higher the uptake rate is for a given substance, the more it is bioaccumulated by organisms. Conversely, an increase in parameter 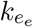 leads to a lower internal concentration (*e.g.*, blue curve in Figure 1.b), consistent with the known underlying biological mechanism: the faster a contaminant is eliminated, the quicker its concentration decreases in organisms.

### 4.2. A one-compartment TK model with several exposure routes, no metabolites

Equations (3) and (15) are simulated for a freshwater shrimp exposed to an organic chlorine compound by contaminated sediment and food. Only natural excretion is considered as elimination process [21] (Figure 2 and Table 2). The corresponding inputs are the exposure concentration via sediment *c_s_* and via food *c_f_* (referring to *c_i_* and *I* = 2), the two uptake rates from sediment and food, 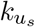 and 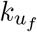 respectively (referring to 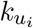 and *I* = 2) and the excretion rate 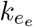 (referring to 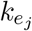 and *J* = 1). When parameter 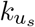 increases, internal concentrations are higher (*e.g.*, blue curve in Figure 2.a) than with the original value (orange curve in Figure 2.a). Biologically speaking, the higher the uptake rate from sediment is for a given substance, the stronger it is bioaccumulated by organisms. Regarding the exposure from food, as parameter 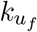 is low in this case, its variation does not influence a lot the internal concentration of the contaminant (Figure 2.b). Biologically speaking, this means that the exposure via sediment is the major route of contamination for these organisms. Besides, as previously shown in section 4.1, an increase in parameter 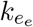 leads to a faster decreasing concentration (*e.g.*, blue curve in Figure 2.c).

**Figure 2:**
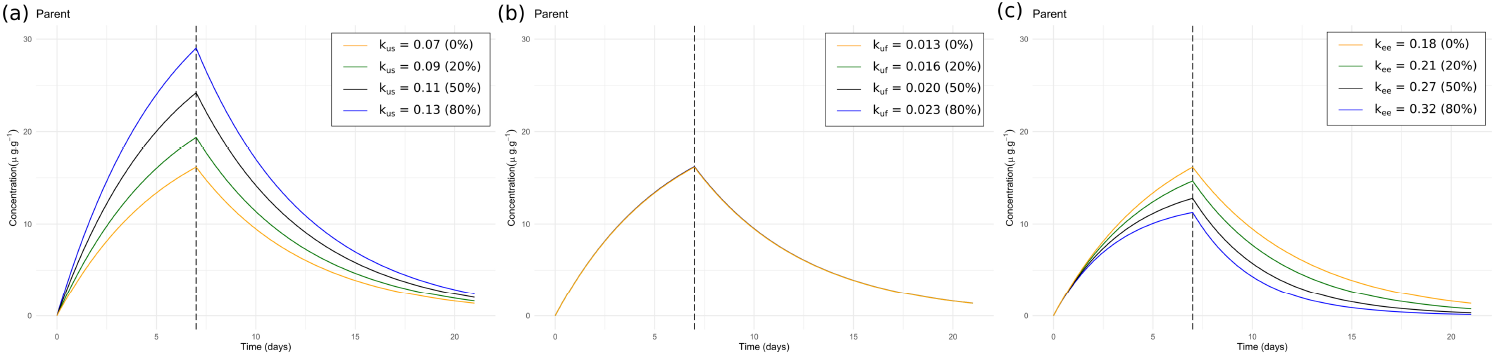
Example of model simulations for a TK model with multiple exposure routes (by sediment and food) and influence of variations of parameters (a) 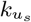 (uptake rate from sediment), (b) 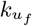 (uptake rate from food) and (c) 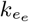 (excretion rate).

### 4.3. A one-compartment TK model with one exposure route, several metabolites

Equations (3), (10), (15) and (23) are simulated for a freshwater shrimp exposed to an organic biocide by contaminated water. Three metabolites derived from the parent compound are considered [1] together with the natural excretion (Figures 3 to 5 and Table 2). The corresponding inputs are the exposure concentration via water *c_w_* (referring to *c_i_* and *I* = 1), the up-take rate 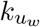 (referring to 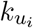 and *I* = 1), the excretion rate 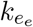 (referring to 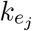 and *J* = 1), the three metabolization rates 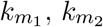 and 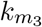 (referring to 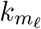 and *L* = 3) and the three elimination rates of the metabolites 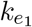, 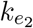 and 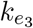 (referring to 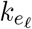 and *L* = 3). When parameter 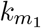 increases, internal concentrations are higher for metabolite 1 (*e.g.*, blue curve in Figure 3.b) than with the original value (orange curve in Figure 3.b). Conversely, internal concentrations are lower for the parent compound (*e.g.*, blue curve in Figure 3.a). Biologically speaking, the more the biotransformation rate for a given metabolite is increasing, the higher is its concentration within organisms due to the highly biotransformation of the parent compound. This leads to a lower internal concentration of the parent compound than with the original value of 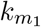 (*e.g.*, blue curve, Figure 3.a). An increase in 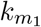 also induces a decrease in the internal concentrations of the other metabolites (Figure 3.c and d). Besides, when parameter 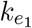 increases, this only affects the internal concentration of metabolite 1 (Figure 4.b). In addition, as previously viewed (sections 4.1 and 4.2), an increase in parameter 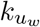 will induce high internal concentrations for both the parent compound and its metabolites (Figure 5). Indeed, the more the internal concentration of the parent compound is increasing, the more the biotransformation process will intensify, leading to high internal concentrations for each metabolite.

**Figure 3:**
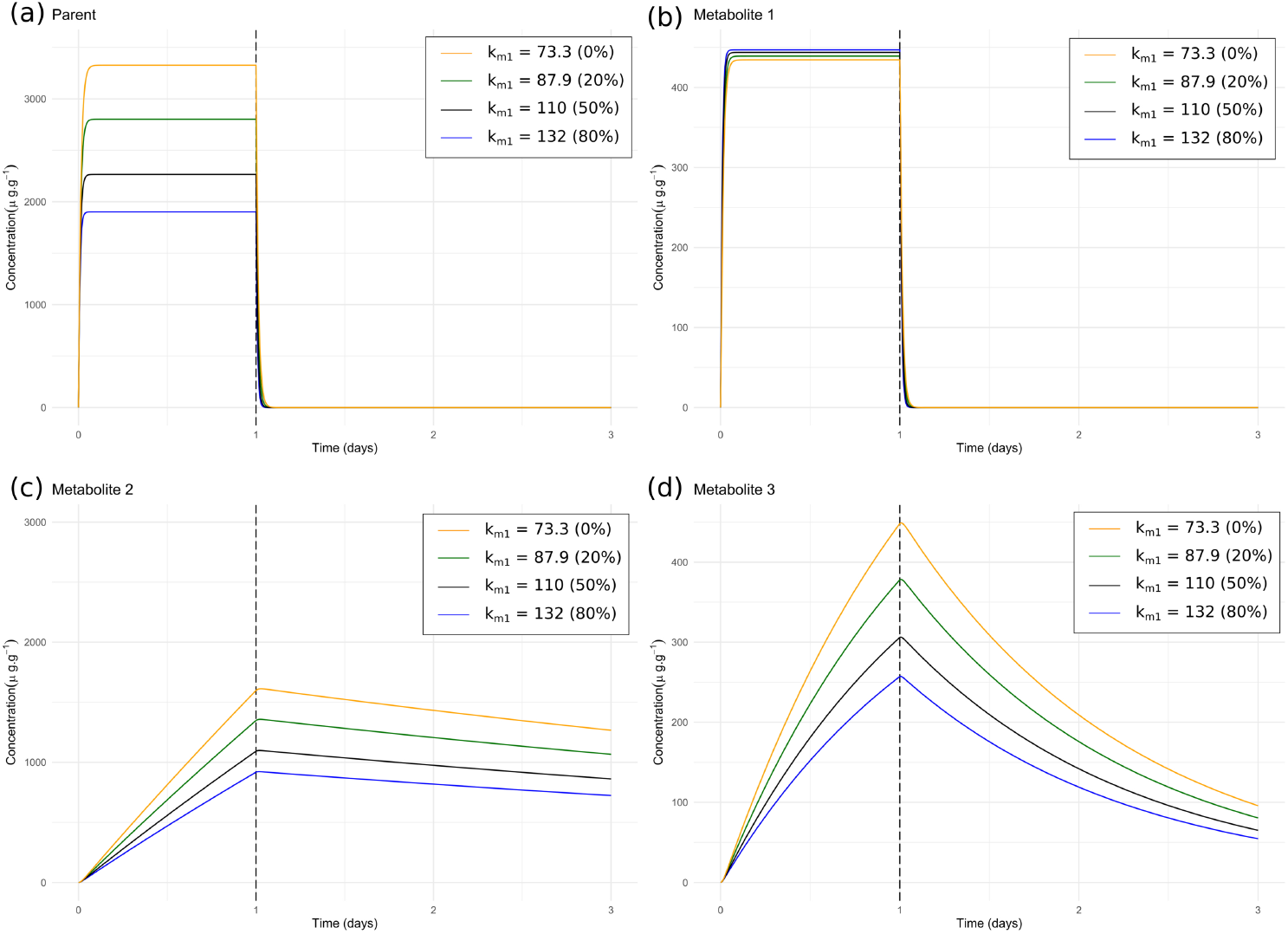
Example of model simulations for a TK model with biotransformation (three metabolites) and influence of the variations of parameter 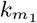 (biotransformation rate for metabolite 1) on (a) the parent compound, (b) metabolite 1, (c) metabolite 2, and (d) metabolite 3.

**Figure 4:**
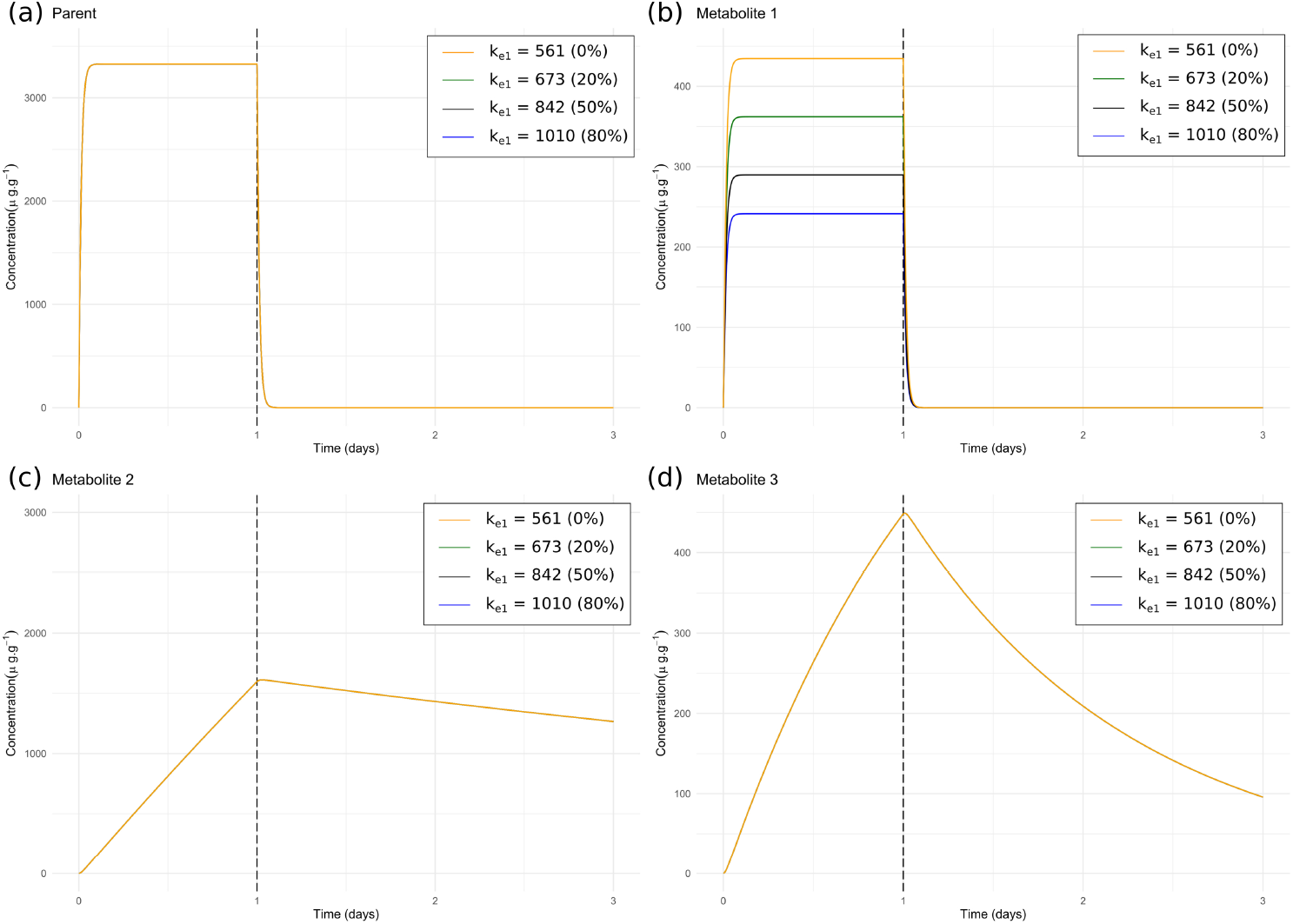
Example of model simulations for a TK model with biotransformation (three metabolites) and influence of the variations of parameter 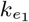 (elimination rate of metabolite 1) on (a) the parent compound, (b) metabolite 1, (c) metabolite 2, and (d) metabolite 3.

**Figure 5:**
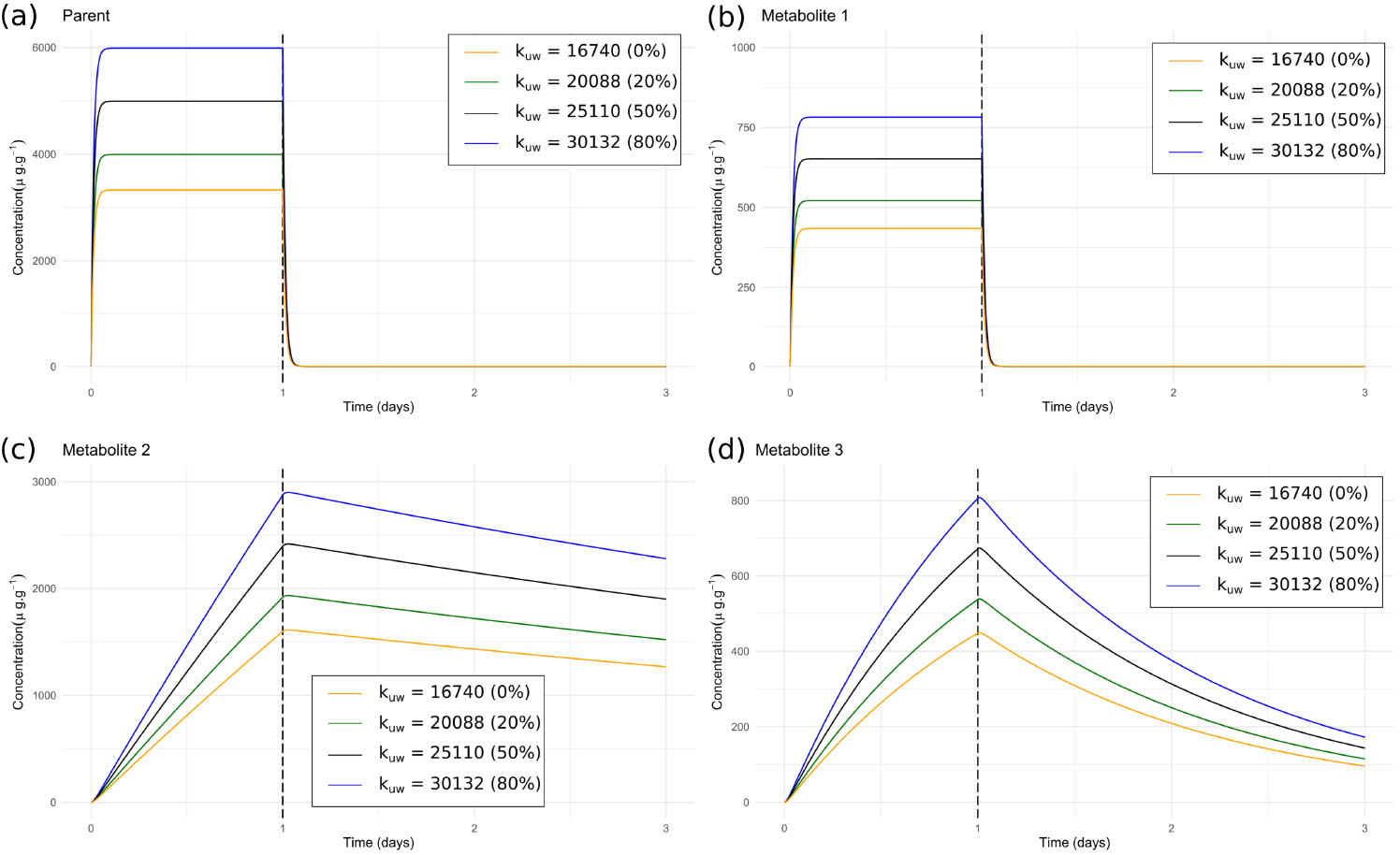
Example of model simulations for a TK model with biotransformation (three metabolites) and influence of the variations of parameter 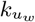 on (a) the parent compound, (b) metabolite 1, (c) metabolite 2, and (d) metabolite 3.

## 5. Conclusion

Based on a full mathematical and exact solving of a generic one-comaprtment TK model, our paper illlustrates its practical use to deal with different kind of species-compound comùbinations of interest. Such a model is today required by regulatory bodies to perform risk assessment of chemical substances. The generic feature of this model is thus in full compliance with the numerous situations with which regulators are expected to be faced. The associated ready-to-use tools we developed to perform different actions such as calibration on experimental data, validation from simulations compared to observed data, or predictions under untested situations, should thus be of valuable help in support of their daily work.

## 6. Acknowledgments

The authors would like to express their sincere thanks to Miléna Kaag, Yacout Lahlou and Nino Molin who designed the MOSAIC*_bioacc_* prediction tool as part of their 4*^th^* year study project at the National Institute of Applied Sciences (INSA) in Lyon (France). This work was made under the umbrella of the French GDR “Aquatic Ecotoxicology” framework aiming at fostering stimulating scientific discussions and collaborations for more integrative approaches. This work was financially supported by the Graduate School H2O’Lyon (ANR-17-EURE-0018) and “Université de Lyon” (UdL), as part of the program “Investissements d’Avenir” run by “Agence Nationale de la Recherche” (ANR).

